# JQ-1 ameliorates schistosomiasis liver fibrosis by suppressing JAK2 and STAT3 activation

**DOI:** 10.1101/2020.08.04.235812

**Authors:** Han Ding, Jiaming Tian, Xuhan Yang, Yongsheng Ji, Saeed El-Ashram, Xin Hou, Cuiping Ren, Jijia Shen, Miao Liu

## Abstract

Schistosomiasis is a serious parasitic infection caused by *Schistosoma*. The parasite deposits eggs in the host liver, causing inflammation that activates hepatic stellate cells (HSCs), which leads to liver fibrosis. Currently, there is no effective therapy for liver fibrosis; thus, treatments are urgently needed. Therefore, in the present study, mice infected with *Schistosoma japonicum* were treated with JQ-1, a small-molecule bromodomain inhibitor with reliable anti-tumor and anti-inflammatory activities. The fibrotic area of the liver measured by computer-assisted morphometric analysis and the expression levels of the cytoskeletal protein alpha smooth muscle actin (α-SMA) and of collagen assessed by quantitative PCR and immunohistochemistry were significantly decreased in the liver following JQ-1 treatment compared with vehicle-treated controls. Total RNA was extracted from the liver of JQ-1–treated *Schistosoma*-infected mice for RNA-sequencing analysis. Gene ontology and Kyoto Encyclopedia of Genes and Genomes analyses indicated that JQ-1 affected biological processes and the expression of cellular components known to play key roles in HSC transdifferentiation into myofibroblasts. In vitro treatment with JQ-1 of JS-1 cells, a mouse HSC line, indicated that JQ-1 significantly inhibited JS-1 proliferation but had no effect on JS-1 activity, senescence, or apoptosis. Western blot results showed that JQ-1 inhibited the expression of phosphorylated JAK2 and phosphorylated STAT3 without altering expression levels of these non-phosphorylated proteins. Taken together, these findings suggested that JQ-1 treatment ameliorated *S. japonicum* egg–induced liver fibrosis, at least in part, by suppressing HSC activation and proliferation through the inhibition of JAK2/STAT3 signaling. These results lay a foundation for the development of novel approaches to treat and control liver fibrosis caused by *S. japonicum*.

**Author summary:** When a host is infected with *Schistosoma,* a common parasite that affects more than a million people and hundreds of thousands of livestock, the parasite deposits eggs in the liver of the host. The eggs lead to liver inflammation, which activates stellate cells in the liver. These stellate cells generate most of the excessive extracellular matrix that replaces healthy liver parenchyma with fibrous tissue, causing liver fibrosis. *Schistosoma japonicum* causes the most severe liver damage of all the schistosome parasites. Therefore, inhibiting the activation of the liver stellate cells and removing the activated stellate cells are key strategies for treating liver fibrosis caused by this parasite. JQ-1 is a potent inhibitor of the BET family of bromodomain proteins and is structurally similar to inhibitors being tested in clinical trials for various types of cancers. Here we found that administering JQ-1 to mice infected with *S. japonicum* decreased the degree of liver fibrosis. JQ-1 inhibited the activation and proliferation of liver stellate cells by blocking the phosphorylation and thus the activation of JAK2 and STAT3 to achieve its therapeutic effects. Thus, this study provides insights into the development of new therapeutic strategies for *Schistosoma*-induced liver fibrosis.

## 1. Introduction

Among the three major schistosomes known to infect humans (*Schistosoma japonicum*, *Schistosoma mansoni*, and *Schistosoma haematobium*), *S. japonicum* causes the most severe pathological damage. In China, *S. japonicum* is a common parasite associated with chronic liver fibrosis [1]. Currently, zoonotic schistosomiasis caused by *S. japonicum* is major public health threat affecting more than a million people and hundreds of thousands of livestock in China [1]. Egg-induced lesions in the liver and intestinal wall of the host are important complications of infection with *S. japonicum*. The immune response of the host to the eggs results in granuloma and fibrosis [2]. Because the mechanism of fibrosis is not fully understood, there is no effective therapy for liver fibrosis. Although the U.S. Food and Drug Administration (FDA) has recently approved pirfenidone and nintedanib as treatments, additional new therapies are needed for treating liver fibrosis [3].

The bromodomain and extra-terminal (BET) family contains four members (bromodomain motif-containing proteins BRD2, BRD3, BRD4, and BRDT) that specifically recognize acetylated lysine residue sites and participate in the regulation of epigenetic protein expression, which plays a key role in regulating various biological processes, including inflammation, the cell cycle, maintenance of the higher-order chromatin structure, and identification of DNA damage signals [4–5]. JQ-1 is a selective inhibitor of the BET family and has been shown to have promising anti-tumor and anti-inflammation effects [6]. JQ-1 has been found to inhibit the development of fibrosis in both a mouse model of hepatic fibrosis induced by carbon tetrachloride (CCl4) and a mouse model of bleomycin-induced pulmonary fibrosis [7–8]. Therefore, we hypothesized that JQ-1 would be effective in the treatment of hepatic fibrosis caused by *S. japonicum*. However, little is known about the mechanisms underlying the effect of JQ-1 on hepatic fibrosis caused by *S. japonicum*.

Therefore, the aim of the present study was to explore the mechanisms of action underpinning the therapeutic effect of JQ-1 by using a mouse model of hepatic fibrosis naturally caused by *S. japonicum*. We found that the degree of liver fibrosis in mice treated with JQ-1 was significantly decreased, mainly through inhibiting the activation and proliferation of hepatic stellate cells (HSCs). The activation and proliferation of HSCs were inhibited by decreasing the phosphorylation and thus activation of Janus kinase 2 (JAK2) and signal transducer and activator of transcription 3 (STAT3). These findings lay a foundation for the development of new approaches for the treatment and control of liver fibrosis caused by *S. japonicum*.

## 2. Materials and methods

### 2.1. Ethics statement

All animal experiments were approved by the Institutional Animal Care and Use Committee at Anhui Medical University (Approval Number: LLSC20140061) and conformed to the guidelines outlined in the Guide for the Care and Use of Laboratory Animals. All infection was performed under anesthesia, and all efforts were made to minimize suffering.

### 2.2. Mouse model of fibrosis, cell culture, and reagents

Female C57BL/6 mice (6 weeks old) were purchased from the Experimental Animal Center of Anhui Province in Hefei, China. Mice were housed under specific pathogen-free conditions at Anhui Medical University, with free access to food and water food. The protocols for the animal experiments were approved by the Institutional Animal Care and Use Committee at Anhui Medical University. Freshwater *Oncomelania hupensis* snails infected with *S. japonicum* (provided by Hunan Provincial Institute of Parasitic Diseases in China) were exposed to light for 3–4 h (25–28 °C) to induce the release of cercariae. Hepatic fibrosis was experimentally induced in 18 mice by infecting them with 18 ± 2 of these cercariae through the skin of abdomen. The infected mice were randomly divided into two groups, experimental and control, with nine mice in each group.

Six weeks after the infection, the mice were treated with praziquantel by intragastric gavage (500 mg/kg/d for 2 days). In the seventh week, mice in the experimental group were injected intraperitoneally (I.P.) with JQ-1 (50 mg/kg body weight/day; Cat. No. HY-13030, MedChem Express; USA), and mice in the control group were injected I.P. with vehicle, that is, (2-hydroxypropyl)-β-cyclodextrin (HP-β-CD; Cat. No. 778966, Sigma; USA) 10% (wt/vol), once daily for 15 days. Animals were humanely killed 24 h after the final injection. They serum and the liver from each mouse were collected for subsequent experimental analyses.

The murine HSC line, JS-1 cells, had been previously purchased from BeNa Culture Collection (Beijing, China) and preserved in our laboratory. For the present study, JS-1 cells were cultured in Dulbecco’s modified Eagle medium with 10% fetal bovine serum.

### 2.3. Egg count in liver tissue

Potassium hydroxide (10%; 1 mL) was added to 0.1 g of liver tissue and digested at 37°C for 2 h. The number of eggs in the sample was then counted using a light microscope.

### 2.4. Assessment of liver fibrosis

Sections (5 μm thick) of formalin-fixed, paraffin-embedded liver specimens were stained with Sirius red. The red-stained collagen fibers were quantified by determining the percentage of the stained area per total liver section as measured by computer-assisted morphometric analysis. Immunostaining for alpha smooth muscle actin (α-SMA), a marker for a subset of activated fibrogenic cells and myofibroblasts, and collagen Ia1 (Col1a1) was performed using polyclonal α-SMA at a dilution of 1:500 and Col1a1 (1:250) primary antibodies (Bioss Inc.; Beijing, China). Immunohistochemical analysis was performed with the PowerVision two-step detection system (Zhongshan Biotechnology Co.; Beijing, China). Six to ten photomicrographs per mouse liver were captured using an inverted microscope (Nikon 80I, Japan). Quantitative and qualitative changes were analyzed using morphometric software (Image-Pro Plus; Media Cybernetics). At least three noncontinuous tissue sections were measured, and the mean values obtained from nine mice were used for statistical analysis.

Hydroxyproline content has been described as a surrogate for collagen content in fibrogenesis. We measured the hydroxyproline content of the liver using a commercial kit (Nanjing Jiancheng Bioengineering Institute; China). Briefly, 1 mL of hydrolysate was added to a liver tissue sample (0.1 g), mixed, and fermented/heated in a boiling water bath for 20 min. The suspension was mixed every few minutes to hydrolyze the tissue components. The resulting components were used to detect the hydroxyproline content according to the kit manufacturer’s instructions. Absorbance was read at a wavelength of 550 nm, and the total hydroxyproline content in the samples was extrapolated from a standard curve.

### 2.5. Serum liver enzyme quantification

The levels of serum alanine aminotransferase (ALT) and aspartate aminotransferase (AST) were determined using a serum aminotransferase test kit (Nanjing Jiancheng Bioengineering Institute; China) according to the manufacturer’s instructions and reported in terms of units per liter.

### 2.6. RNA sequencing and data analysis

Three individual liver samples from each group were pooled to generate one sample for RNA sequencing. Liver samples were finely ground, and total RNA was extracted using TRIzol (Invitrogen; Carlsbad, CA, USA). The quantity and purity of the total RNA were assessed using a Bioanalyzer 2100 system. A 5-μg sample of total RNA was used to isolate poly(A) mRNA with a poly(T) oligo primer attached to magnetic beads (Invitrogen). Briefly, mRNA was extracted from total RNA using oligo (dT) magnetic beads and sheared into short fragments. These fragments of mRNA were then used as templates for cDNA synthesis. The cDNAs were amplified by polymerase chain reaction (PCR) to complete the library. The cDNA library was constructed from the RNA samples and sequenced with an Illumina Hiseq 2500 system according to the manufacturer’s instructions. Differentially expressed genes were submitted to enrichment analysis by gene ontology (GO) and Kyoto Encyclopedia of Genes and Genomes (KEGG). *P* values were calculated using the Benjamini-corrected modified Fisher’s exact test, and *P* ≤ 0.05 was considered statistically significant.

### 2.7. Quantitative PCR

Total RNA was extracted from the liver of each mouse. The mRNA was reverse transcribed to cDNA using a PrimeScript RT reagent kit (Takara; Dalian, China). Real-time reverse transcription quantitative PCR (qPCR) was performed in duplicate using 1 μL of cDNA from each sample and SYBR Premix Ex Taq 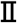 (Takara) according to the manufacturer’s instructions. All reactions were performed using an ABI-Prism StepOnePlus sequence detector system under the following conditions: 10 s at 95 °C; 40 cycles of 15 s at 95 °C and 40 s at 60 °C followed by 15 s at 95 °C; 1 min at 60 °C; and 95 °C for 15 s. This allowed for the generation of a standard regression curve using threshold settings and software analysis. Primers for qPCR were designed and synthesized by Sangon Biotech Co. Ltd. Quantification was performed by comparing the cycle threshold (Ct) values of each sample after normalization to GAPDH. Primer sequences are summarized in Supplementary Table 1. The relative quantification levels were calculated using the 2^(−ΔΔCT)^ method.

### 2.8. Cell viability assay

The mouse HSC line JS-1 cells were cultured on six-well chamber slides in the presence of DMSO (0.1%) or JQ-1 (500 nM) for 24 h. The exponentially growing cells were trypsinized and washed twice with phosphate-buffered saline (PBS). Then, 450 μL of cell suspension was mixed with 50 μL of 0.4% trypan blue dye and left for 5 min at room temperature. Trypan blue exclusion was assessed, viable cells were counted using a hemocytometer, and the percentage of viable cells was determined.

### 2.9. Cell proliferation assay

JS-1 cells were cultured on six-well chamber slides in the presence of DMSO (0.1%) or JQ-1 (500 nM) for 24 h. The Cell Counting Kit-8 (CCK-8; Dojindo Laboratories; Japan) solution (10 μL/well) was added to measure cell proliferation at specific intervals (0, 12, and 24 h of incubation). After 2 h of further incubation at 37 °C in 5% CO_2_, the absorbance of each well was measured at a wavelength of 450 nm by a Model 680 microplate reader (Bio-Rad; Richmond, CA, USA).

### 2.10. Terminal deoxynucleotidyl transferase dUTP nick end labeling (TUNEL) Assay

JS-1 cells were cultured on six-well chamber slides in the presence of DMSO (0.1%) or JQ-1 (500 nM) for 24 h. After incubation, JQ-1–induced apoptosis of JS-1 cells was determined using a One-Step TUNEL Apoptosis Assay Kit (Beyotime; China) according to the manufacturer’s instructions. Briefly, JQ-1–treated cells were washed twice with ice-cold PBS, fixed with 4% paraformaldehyde for 30 min, and permeabilized with 0.5% Triton X-100 for 5 min. The cells were subsequently incubated with the TUNEL detection reagent at 37 °C for 1 h. Cell fluorescence was observed under a fluorescence microscope.

### 2.11. Senescent cell staining

JS-1 cells cultured on six-well chamber slides in the presence of DMSO (0.1%) or JQ-1 (500 nM) were stained for a senescence-associated marker using a β-galactosidase staining kit (Beyotime; Shanghai, China).

### 2.12. Western blot analysis

JS-1 cells were rinsed with ice-cold PBS, harvested, and lysed in a lysis buffer (HEPES, 25 mM; 1.5% Triton X-100; 1% sodium deoxycholate; 0.1% sodium dodecyl sulfate [SDS]; NaCl, 0.5 M; EDTA, 5 mM; NaF, 50 mM; sodium vanadate, 0.1 mM; phenylmethylsulfonyl fluoride, 1 mM; and leupeptin, 0.1 g/L; pH 7.8) containing a protease inhibitor (Roche Applied Science; Indianapolis, IN, USA). Cell lysates were solubilized in SDS sample buffer, separated by 12% SDS–polyacrylamide gel electrophoresis with 60 μg of protein loaded per lane, and transferred to a nitrocellulose membrane, which was blocked with blocking buffer (5% non-fat dry milk) for 2 h at room temperature. To detect activation of the JAK2-STAT3 signaling pathway, the membranes were then incubated with the primary antibodies against phosphorylated (p)-JAK2 (Cat. No. 3771), JAK2 (Cat. No. 3230), p-STAT3 (Cat No. 4113), STAT3 (Cat. No. 4904), and β-actin (Cat. No. 3700) (all from Cell Signaling Technology; USA), all diluted 1:1000 and incubated overnight at 4 °C. To detect the expression level of α-SMA, collagen I, and collagen III, the membranes were then incubated with the primary antibodies against α-SMA (Cat. No. 3771, Abcam; Cambridge, MA, USA), collagen Ia1 (Col1a1; Servicebio; Wuhan, China), collagen III (Col3a1; Servicebio), and GAPDH (Cat. No. 3700, Abcam) followed by incubation with the corresponding horseradish peroxidase–conjugated secondary antibody (1:3,000; EMD Millipore, MA, USA) for 2 h at room temperature. Finally, the membranes were developed by enhanced chemiluminescence solutions (Thermo Fisher Scientific) and visualized using the chemical luminescence imaging apparatus FluorChem FC3 (GeneTech; Shanghai, China). Immunoreactive bands were quantified using densitometry (multiplex band analysis from FluorChem FC3).

### 2.13. Statistical analysis

All data were obtained in three independent experiments, with triplicate samples used for each experiment. Data sets were analyzed with Student’s *t* test using GraphPad Prism 6.01 software. Data are presented as means ± SEM. Two-sided values of *P* < 0.05 were considered statistically significant.

## 3. Results

### 3.1 JQ-1 ameliorates liver fibrosis caused by *S. japonicum*

In the seventh week after *S. japonicum* infection, mice in the experimental group were injected with JQ-1, and mice in the control group were injected with the vehicle HP-β-CD, once daily for 15 days. All mice were humanely killed after 15 days of treatment (Fig. 1A). The numbers of *S. japonicum* eggs were counted in the livers and were found to be similar in both groups (Fig. 1B). As shown in Fig. 1C, the overall color of the liver from mice in the HP-β-CD group was dark and the surface was irregular, suggestive of severe fibrosis. In comparison, the color of the liver in the JQ-1–treated group was lighter and more vivid, and the surface was relatively smooth.

**Fig. 1.**
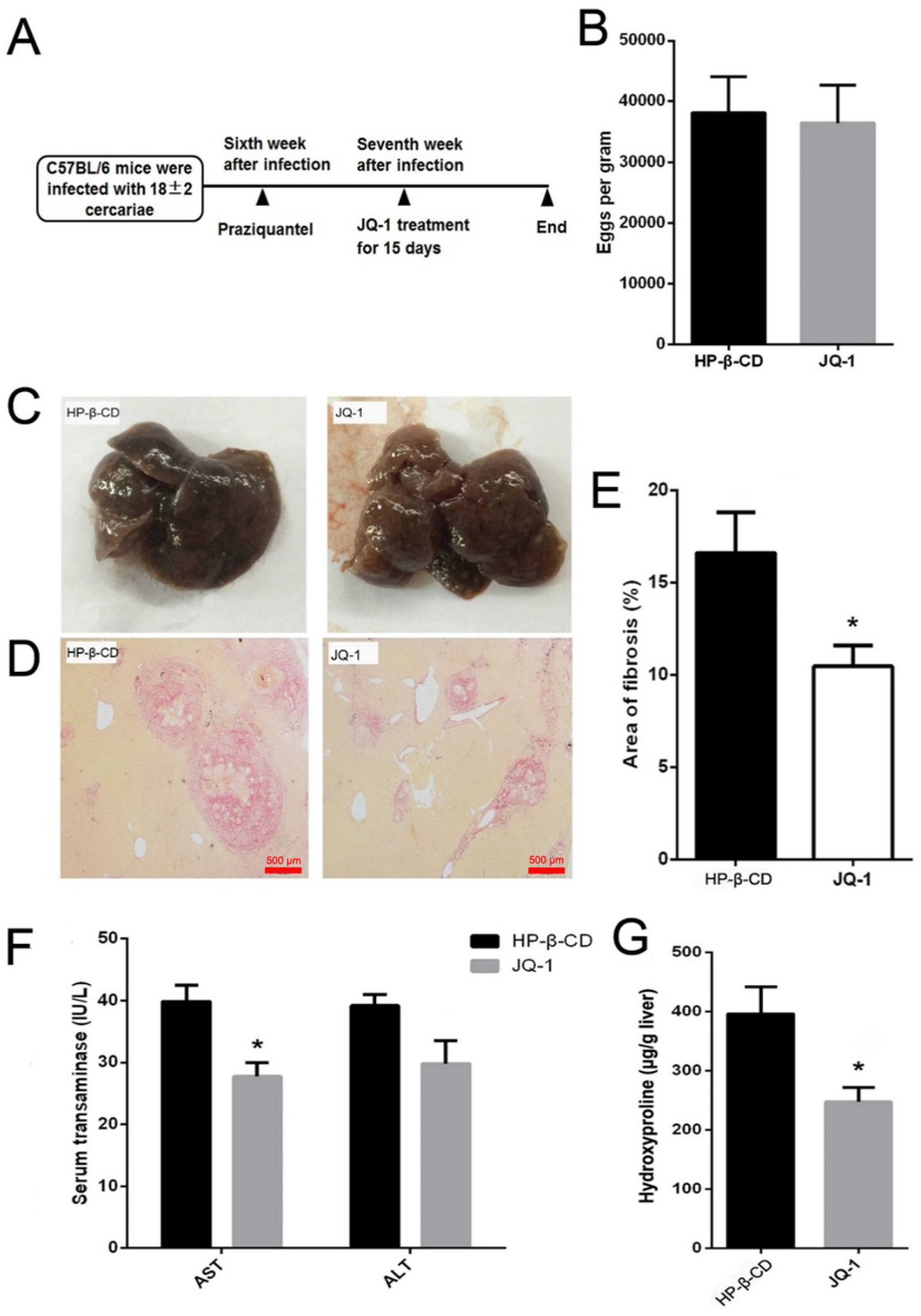
JQ-1 treatment ameliorates liver fibrosis caused by *S. japonicum*. (A) Therapeutic regimen used in the mouse model of liver fibrosis. (B) Effect of JQ-1 or vehicle (HP-β-CD) treatment on the number of eggs per gram in the liver. (C) Effect of JQ-1 treatment on general liver morphology of mice infected with *S. japonicum*. (D) Effect of JQ-1 treatment on hepatic fibrosis in mice infected with *S. japonicum*. (E) Area of the liver with hepatic fibrosis as measured by computer-assisted morphometric analysis. (F) Effect of JQ-1 treatment on serum alanine aminotransferase (ALT) and aspartate aminotransferase (AST) levels and (G) liver hydroxyproline levels in mice infected with *S. japonicum*. Data represent the mean ± SEM (n = 9 for each group). Asterisks denote statistically significant differences (Student’s *t* test, *P* < 0.05) vs. the HP-β-CD–treated control group.

Three non-continuous fixed, embedded, and sectioned liver slices were selected from nine mice for Sirius red staining and morphological analysis. As shown in Fig. 1D and 1E, Sirius red staining indicated that the area of liver fibrosis in the group treated with JQ-1 was significantly smaller than that in the control HP-β-CD group. Fibrosis was observed throughout the liver of mice in the HP-β-CD group as wide scars. By contrast, the infiltration of inflammatory cells and collagen fiber in the liver tissue was significantly reduced in the group of mice treated with JQ-1.

The degree of liver cell damage was evaluated by measuring the enzyme levels of AST and ALT in the serum of the mice. As shown in Figure 1F, serum transaminase activity in the JQ-1 group was significantly lower than that in the HP-β-CD group (*P* < 0.05). In the JQ-1 group, the level of hydroxyproline in the liver tissue of mice was also significantly lower than that in the HP-β-CD group (Figure 1G, *P* < 0.05). Taken together, these results indicated that the degree of liver fibrosis in the JQ-1 group was significantly decreased.

### 3.2. JQ-1 treatment decreases collagen expression in the liver by reducing activation of HSCs

The immunohistochemical results showed that the expression levels of α-SMA and Col1a1 in mouse liver significantly decreased after JQ-1 treatment (Fig. 2A and 2B). The expression level of α-SMA was significantly lower in the JQ-1 treated group than in the control group (*P* < 0.05). In addition, the expression levels of Col1a1 and Col3a1 in the HSCs of the JQ-1–treated group were also significantly lower than those in the control group (Fig. 2C, *P* < 0.05). These results indicated that JQ-1 treatment decreased the expression of collagen in the liver by inhibiting the activation of HSCs in the liver to ameliorate hepatic fibrosis.

**Fig. 2.**
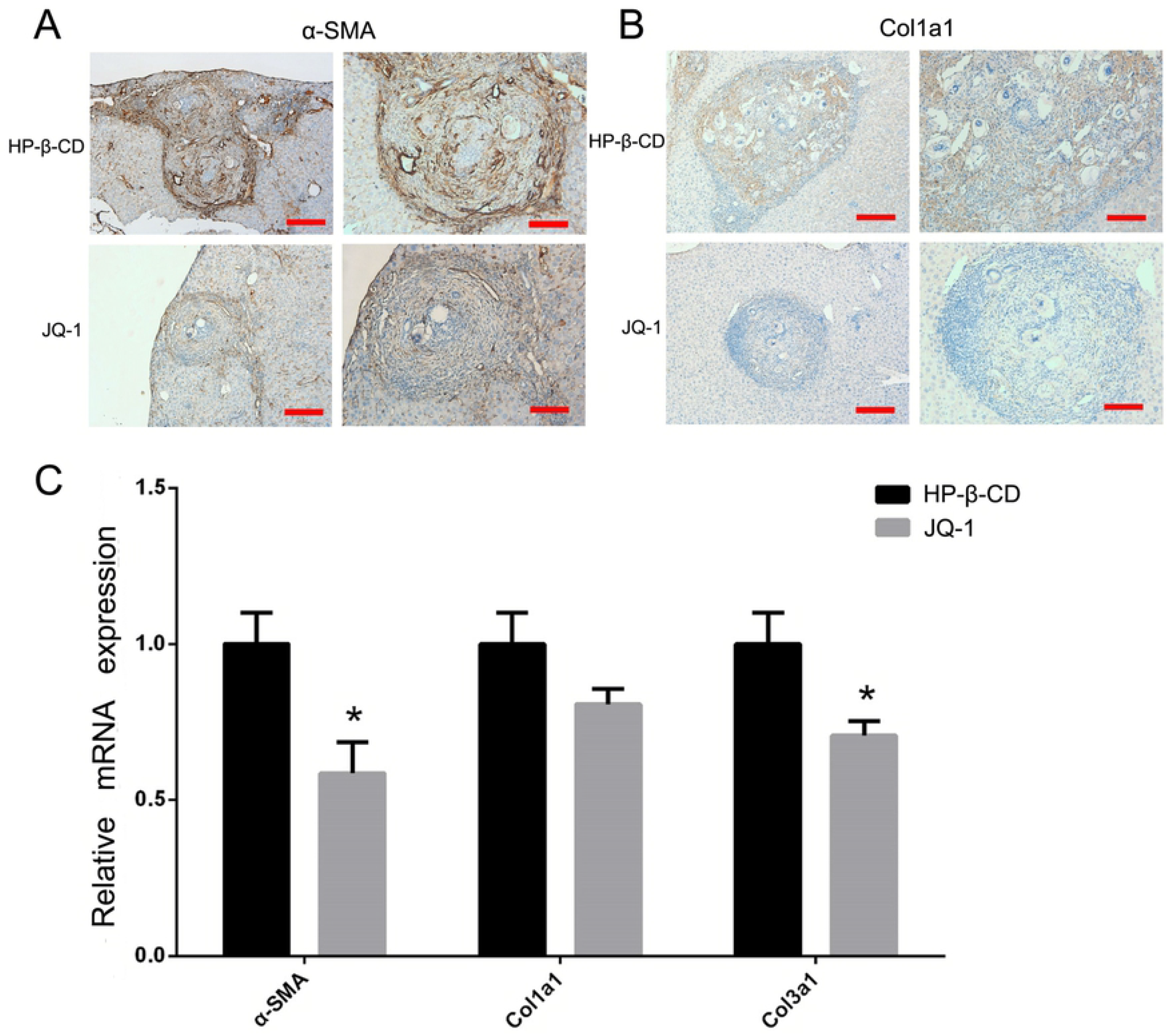
Effect of JQ-1 treatment on expression levels of α-SMA and collagen in mouse liver. Immunohistochemistry images of (A) α-SMA and (B) Col1a1 expression in mouse liver after JQ-1 or vehicle (HP-β-CD) treatment. (C) Quantitative PCR analysis of liver α-SMA and liver collagen protein levels after JQ-1 or vehicle (HP-β-CD) treatment in mice. Data represent the mean ± SEM (n = 9 for each group). each performed in triplicate. Asterisks denote statistically significant differences (Student’s *t* test, \**P* < 0.05).

### 3.3. GO and KEGG pathway enrichment analyses

After RNA-sequencing analysis, we analyzed 25,228 genes (Supplemental Information I). Differential gene clustering analysis results showed that compared with the HP-β-CD–treated control group (Group 1; G1), there were 550 differentially expressed genes (≥2-fold difference) in the JQ-1–treated group(Group 2; G2), of which 263 genes were upregulated and 287 genes were downregulated (*P* < 0.05) (Figure 3A and Supplemental Information II). The results of subsequent analyses using qPCR were consistent with the bioinformatics analyses and identified the differentially expressed genes (Fig. 3B).

**Fig. 3.**
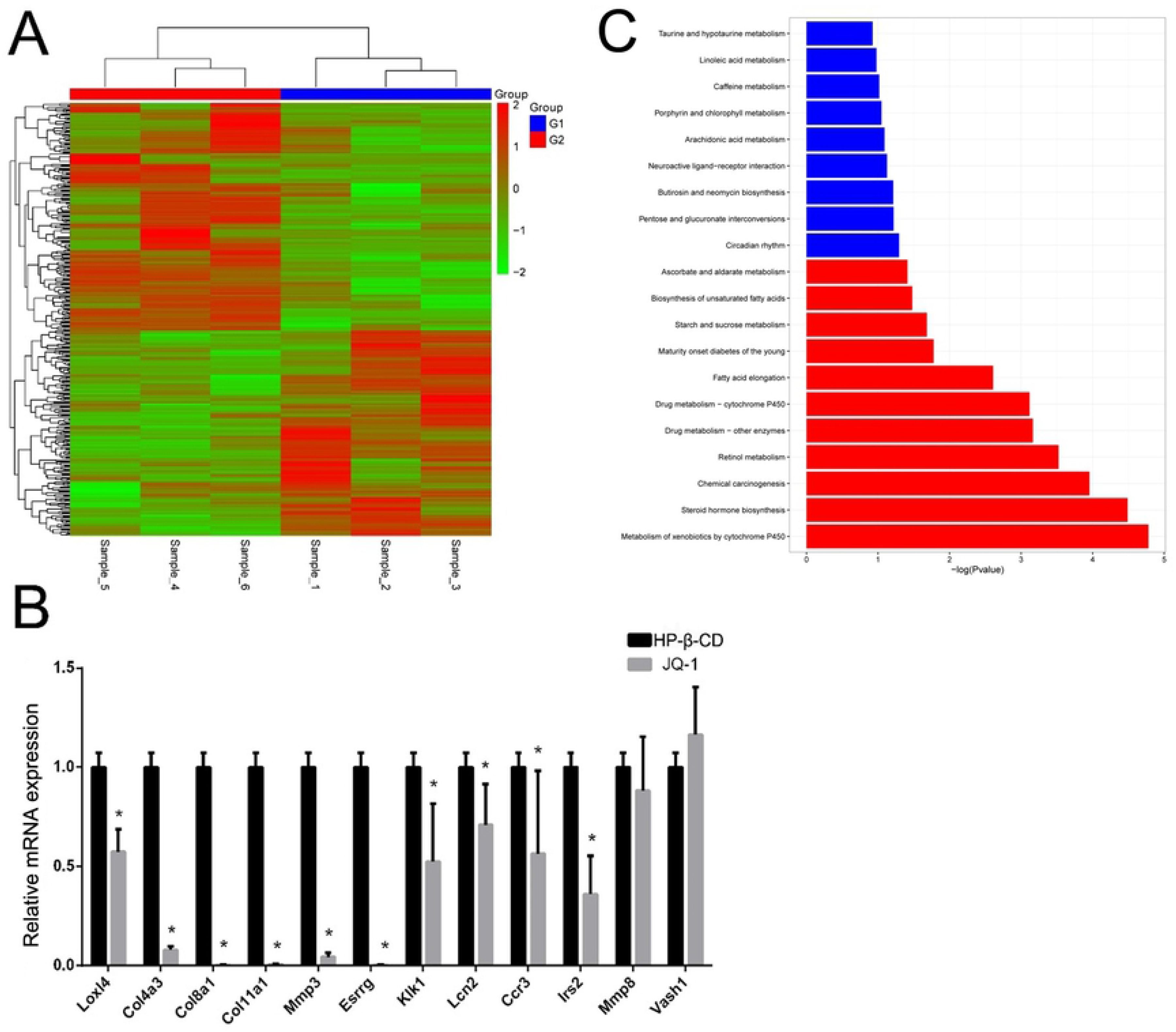
Bioinformatics Analysis. (A) Heat map of differential clustering analysis of mRNA in the JQ-1 treated group (Group 2; G2) versus the HP-β-CD group (Group 1; G1). (B) Differentially expressed genes identified using quantitative PCR. The mRNA levels are expressed relative to those in controls following normalization to GAPDH. Data represent the mean ± SEM (n = 9 for each group). Asterisks denote statistically significant differences (Student’s *t* test, *\*P* < 0.05) vs. HP-β-CD group. (C) KEGG analysis of downregulated genes involved in the metabolic pathway of the proteins. The abscissa indicates *P* values; red bars, highly enriched and meaningful metabolic pathways; and blue bars, a less enriched and reference metabolic pathway. The ordinate shows the name of the metabolic pathway.

The 186 downregulated genes were submitted to the Database for Annotation, Visualization and Integrated Discovery (DAVID) for GO term analysis. The GO annotation indicated that JQ-1 affected a broad spectrum of biological processes and expression of cellular components that are known to play key roles in HSC transdifferentiation into myofibroblasts. These components included the extracellular regions, extracellular matrix, extracellular region part and receptor activity, regulation of ion transport, lipid metabolic processes, and integrin pathway (Supplementary Table 2). Based on the KEGG pathway database, a pathway analysis was performed to predict the significantly enriched metabolic pathways and signal transduction pathways in the host differentially expressed mRNAs. The significantly enriched pathways are shown in Figure 3C. There were six significant signaling pathways: xenobiotic biodegradation and metabolism, lipid metabolism, cancers, metabolism of cofactor and vitamins, endocrine and metabolic diseases, and carbohydrate metabolism (Supplementary Table 3).

### 3.4. Cytology results

In cytology studies, we found that although JQ-1 had no effect on cell viability (Fig. 4A) or senescence-associated β-galactosidase activity (Fig. 4B), it significantly inhibited the proliferation of JS-1 cells (Fig. 4C). The results of a TUNEL assay (Fig. 4D and 4E) provided further evidence that this inhibition of JS-1 cells was specific to proliferation and was not caused by apoptosis.

**Fig. 4.**
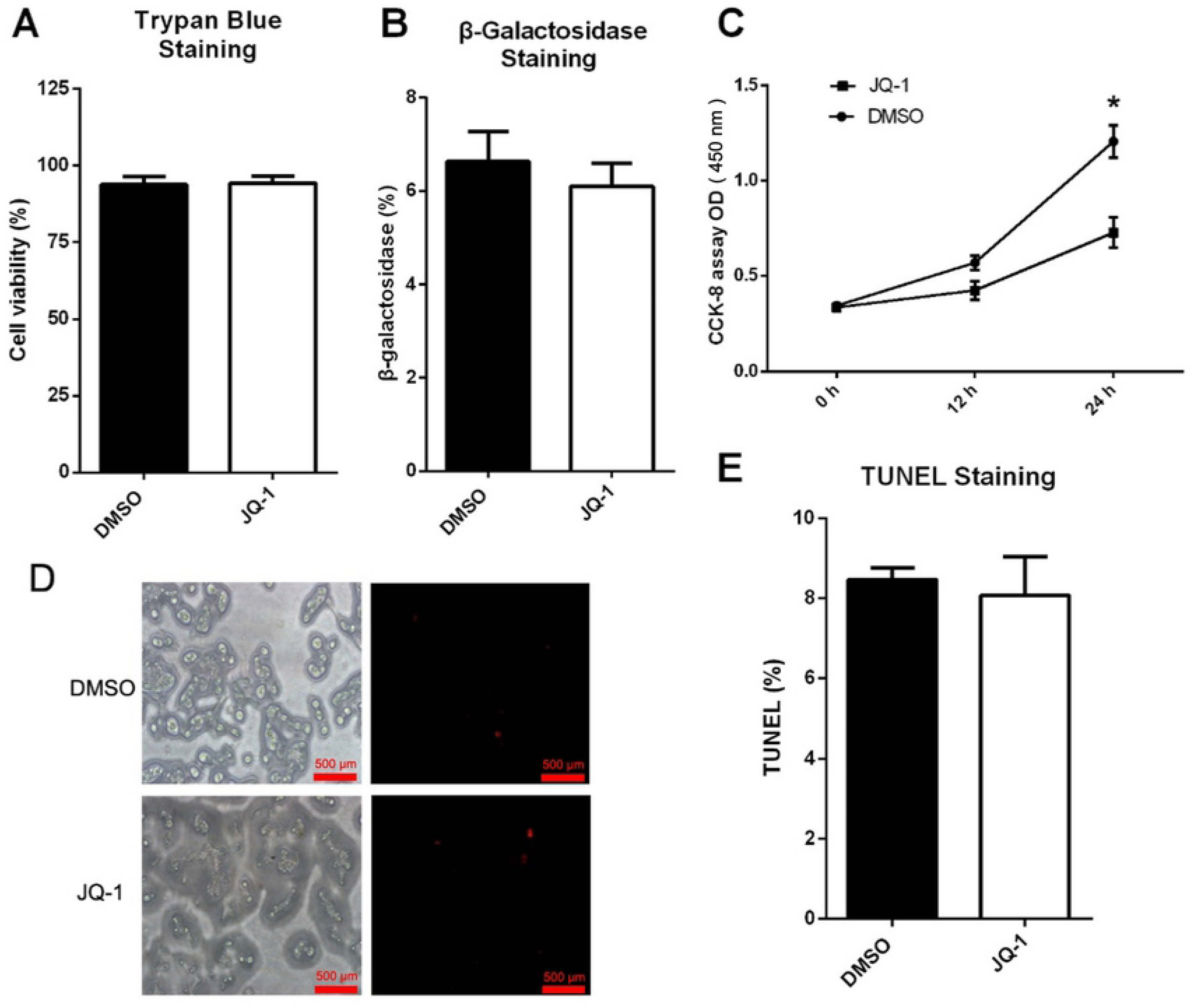
Effect of JQ-1 treatment on hepatic stellate cell (HSC) activity, aging, proliferation, and apoptosis. (A) Cell viability of JQ-1–treated (500 nM) HSCs as measured by trypan blue staining. (B) Cellular senescence of JQ-1–treated (500 nM) HSCs measured by β-galactosidase staining. (C) Cell proliferation of JQ-1–treated (500 nM) HSCs measured by CCK-8 (\**P* < 0.05). OD represents optical density. (D) Apoptosis in JQ-1–treated (500 nM) HSCs measured by a TUNEL assay. Data represent the mean ± SEM of at least three independent experiments, each performed in triplicate. Asterisks denote statistically significant differences (Student’s *t* test, \**P* < 0.05).

### 3.5. JQ-1 inhibits activation of JS-1 cells

We used α-SMA to assess the degree of HSC activation. Using western blot and quantitative PCR analyses, we found that the expression level of α-SMA in JQ-1–treated JS-1 cells was significantly lower than that in the vehicle-treated (DMSO) control group (Fig. 5A; *P* < 0.05). We also found that the protein expression levels of collagen as assessed with Col1a1 and Col3a1 in JQ-1–treated JS-1 cells were significantly lower than those in the vehicle-treated (DMSO) control group (Fig. 5B; *P* < 0.05). These results indicated that JQ-1 treatment decreases the expression of collagen in the liver by inhibiting the activation of JS-1 cells.

**Fig. 5.**
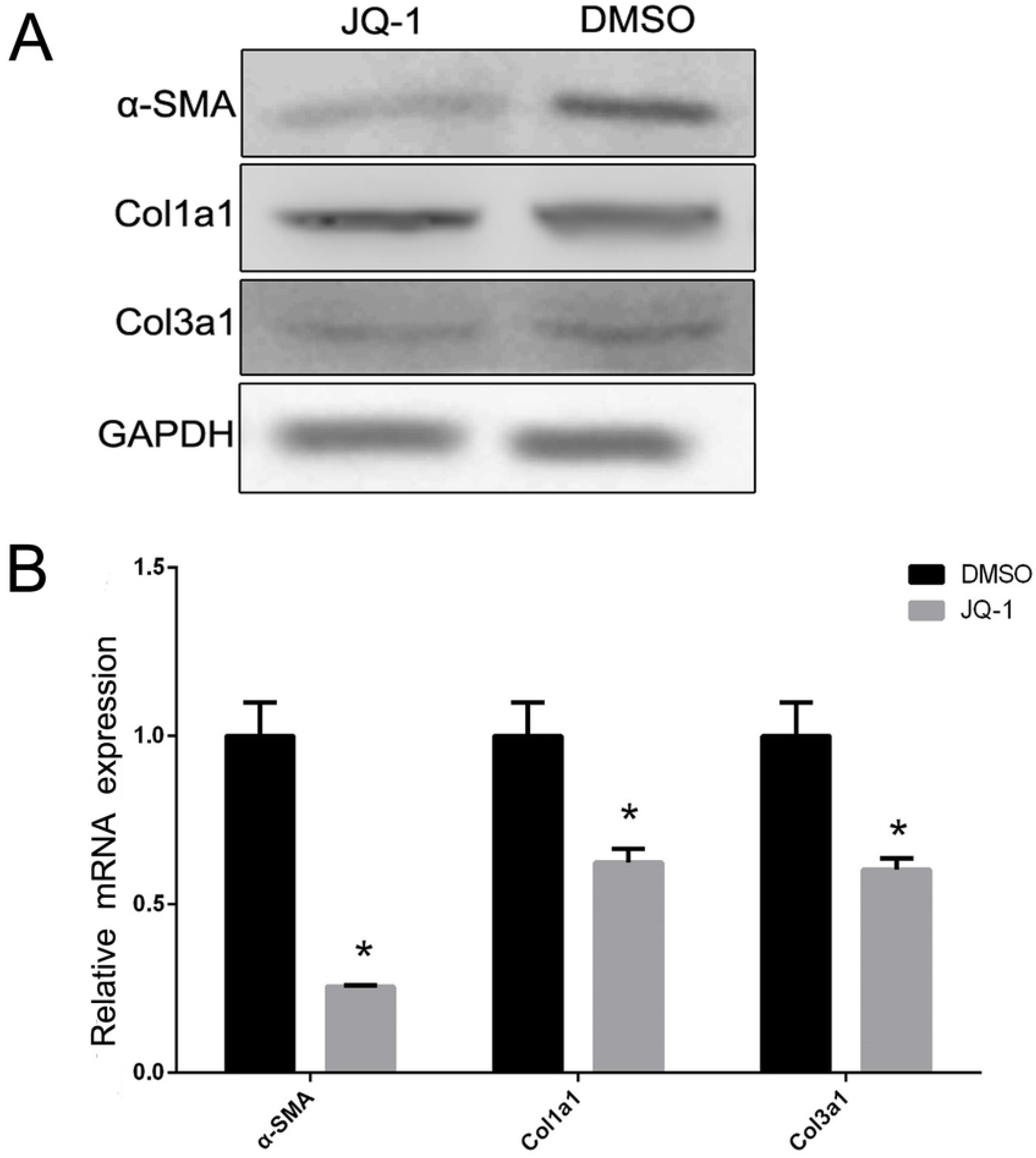
Effect of JQ-1 treatment on the expression levels of α-SMA and collagen in hepatic stellate cells (HSCs). (A) Representative western blot images for α-SMA, Col1a1, and Col3a1 in HSCs after JQ-1 treatment.(B) Relative quantitative PCR analysis of α-SMA and collagen levels in HSCs after JQ-1 treatment. Data represent the mean ± SEM of at least three independent experiments, each performed in triplicate. Asterisks denote statistically significant differences (Student’s *t* test, \**P* < 0.05).

### 3.6. JQ-1 treatment inhibits the phosphorylation of JAK2 and STAT3 in JS-1 cells

As shown in Fig. 6, the expression levels of p-JAK2 and p-STAT3 were significantly decreased in JS-1 cells treated with JQ-1 compared with those in the DMSO-treated (vehicle control) group (*P* < 0.05). These results suggested that JAK2 and STAT3 were less active in JS-1 cells after treatment with JQ-1.

**Fig. 6.**
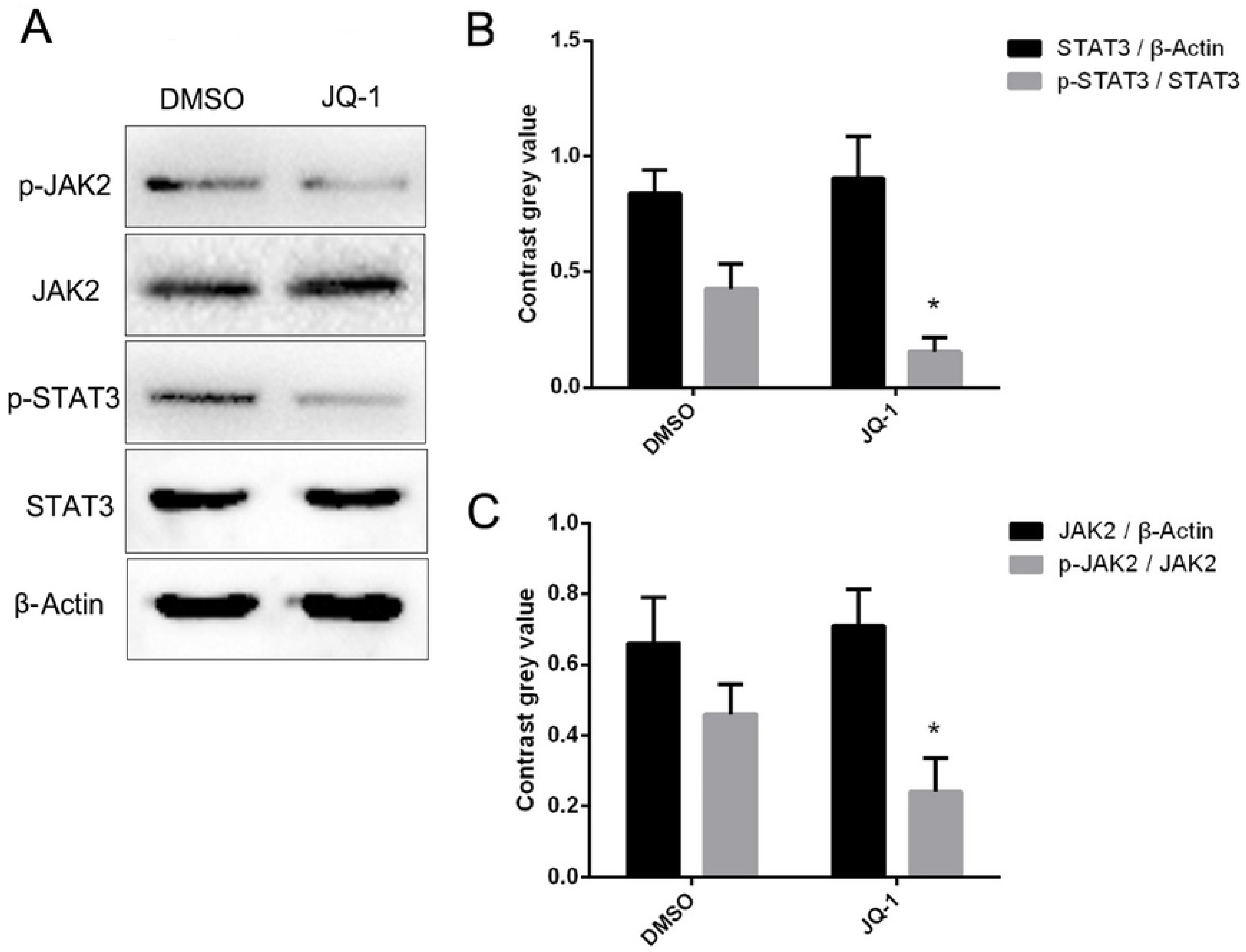
JQ-1 inhibits STAT3 and JAK2 phosphorylation. (A) Representative western blot images for JAK2, phosphorylated (p)-JAK2, STAT3, and p-STAT3 in JS-1 cells after JQ-1 or vehicle control (DMSO) treatment.(B) Relative quantitative analysis of JAK2, p-JAK2, STAT3, and p-STAT3 levels. β-Actin was used as an internal control. Data represent the mean ± SEM of at least three independent experiments, each performed in triplicate. Asterisks denote statistically significant differences (Student’s *t* test, \**P* < 0.05).

## 4. Discussion

Previous studies have shown that JQ-1 blocks the expression of profibrotic factors by competitively binding BRD4 in the carbon tetrachloride (CCl4)-induced model of liver fibrosis in mice, inhibiting the development of fibrosis [7]. However, the observed effect and mechanisms in the CCl4 model may not be applicable to infectious liver fibrosis. In the present study, a mouse model of liver fibrosis was generated by infection with *S. japonicum* to gain insight into the effects and mechanisms underlying treatment with JQ-1 in *S. japonicum* egg–induced liver fibrosis. We found that JQ-1 treatment significantly attenuated *S. japonicum* egg–induced fibrosis and significantly improved the gross appearance of the liver (Fig. 1). We also observed that JQ-1 treatment significantly inhibited expression of α-SMA, a marker of activation of HSCs in liver tissue (Fig. 1).

The process of fibrosis formation from granular tissue induced by *S. japonicum* eggs deposited in the liver is dependent on HSCs, which can undergo activation and transform into myofibroblast-like cells [9]. This activation is characterized by the gradual release of intracellular vitamin A, a high rate of cell proliferation, the synthesis of a fibrogenic matrix rich in type I or type III collagen, activation of α-SMA, and a high expression of extracellular matrix (ECM) [10]. Gradual deposition of ECM leads to structural and functional disorders of the liver [11]. Because HSCs are the main fibrogenic precursor cells in the liver, the blockade, reversal, or both of this activation determines the efficacy of antifibrotic therapy and is key for the treatment of schistosomiasis-induced liver fibrosis [12].

HSCs are typically non-proliferative in normal liver tissue but become activated after liver injury or in vitro culture [13]. Non-activated HSCs are characterized by the storage of vitamin A and the accumulation of liposomes [14]. When HSCs are stimulated and become activated, they are transdifferentiated into myofibroblasts with characteristics of hyperplasia, contraction, inflammation, the disappearance of lipid droplets, chemotaxis, and a massive deposition of ECM [13]. Our sequencing results showed that genes downregulated by JQ-1 treatment and the signaling pathways regulated by these downregulated genes were involved in vitamin metabolism, lipid metabolism, and ECM synthesis. This finding indicates that treatment with JQ-1 decreased the level of vitamin A and lipid metabolism in HSCs, resulting in a decrease in ECM content, and suggests that JQ-1 reduced the severity of liver fibrosis by inhibiting activation of HSCs. Our immunohistochemistry and western blot analysis results showed that the expression levels of collagen I and collagen III also decreased in liver tissue after treatment with JQ-1.

The JAK/STAT signaling pathway transmits extracellular information from the cell membrane to the nucleus to modulate the transcription of target genes, including genes whose protein products are crucial for defense against pathogens, differentiation, proliferation, and apoptosis [15]. The JAK2/STAT3 signaling pathway is directly involved in HSC activation and transdifferentiation and subsequent formation of hepatic fibrosis [16]. The blockade of the JAK2/STAT3 signaling pathway impedes the morphological transdifferentiation of HSCs and reduces mRNA expression of profibrotic genes [17]. Cao et al. (2019) found that intragastric administration of dextran sulfate sodium reduced hepatic fibrosis by inhibiting the JAK2/STAT3 signaling pathway, both in vitro and in vivo [18]. Magnesium isoglycyrrhizinate ameliorates high fructose-induced liver fibrosis in rats via inhibiting the JAK2/STAT3 signaling pathway and TGF-β1/Smad signaling [19].

Recent studies have shown that JQ-1 inhibits the growth of pancreatic cancer cells, regulates the expression of inflammatory cytokines associated with tumor growth, and regulates the expression of several proteins known to be important in pancreatic cancer, such as c-Myc, p-Erk1/2, interleukin 6, and p-STAT3 [20]. Moreover, a study investigating JQ-1 treatment of acute myeloid leukemia cells indicated that this treatment inhibits the expression of p-STAT3 [21]. JQ-1 has been shown to inhibit CCl4-induced liver fibrosis by preventing HSC activation; however, the mechanism underlying the inhibition of HSC activation requires further study [7, 22].

To further investigate the mechanism by which JQ-1 inhibits HSC activation in the present study, the mouse HSC line (JS-1 cells) was cultured and JQ-1 administered in subsequent in vitro experiments. We found that the expression levels of p-JAK2 and p-STAT3 were significantly reduced in the JQ-1 treated JS-1 cells (Fig. 6), suggesting that JQ-1 inhibits activation of HSCs by suppressing the phosphorylation of JAK2 and STAT3. As shown in Fig. 7, JQ-1 inhibits the JAK2/STAT3 signaling pathway in JS-1 cells, thereby blocking the activation and proliferation of HSCs and reducing the severity of liver fibrosis. JQ-1 can inhibit all BET proteins that disturb the binding of BET to acetylated lysine, which can regulate numerous targets to affect hepatic fibrosis. Suppressing JAK2 and STAT3 activation may be a JQ-1 mechanism that inhibits schistosomal hepatic fibrosis. However, our study did not determine whether one of the BET proteins is responsible for the transcription of genes involved in the activation and proliferation of HSCs; thus, the underlying mechanism requires future investigation.

**Fig. 7.**
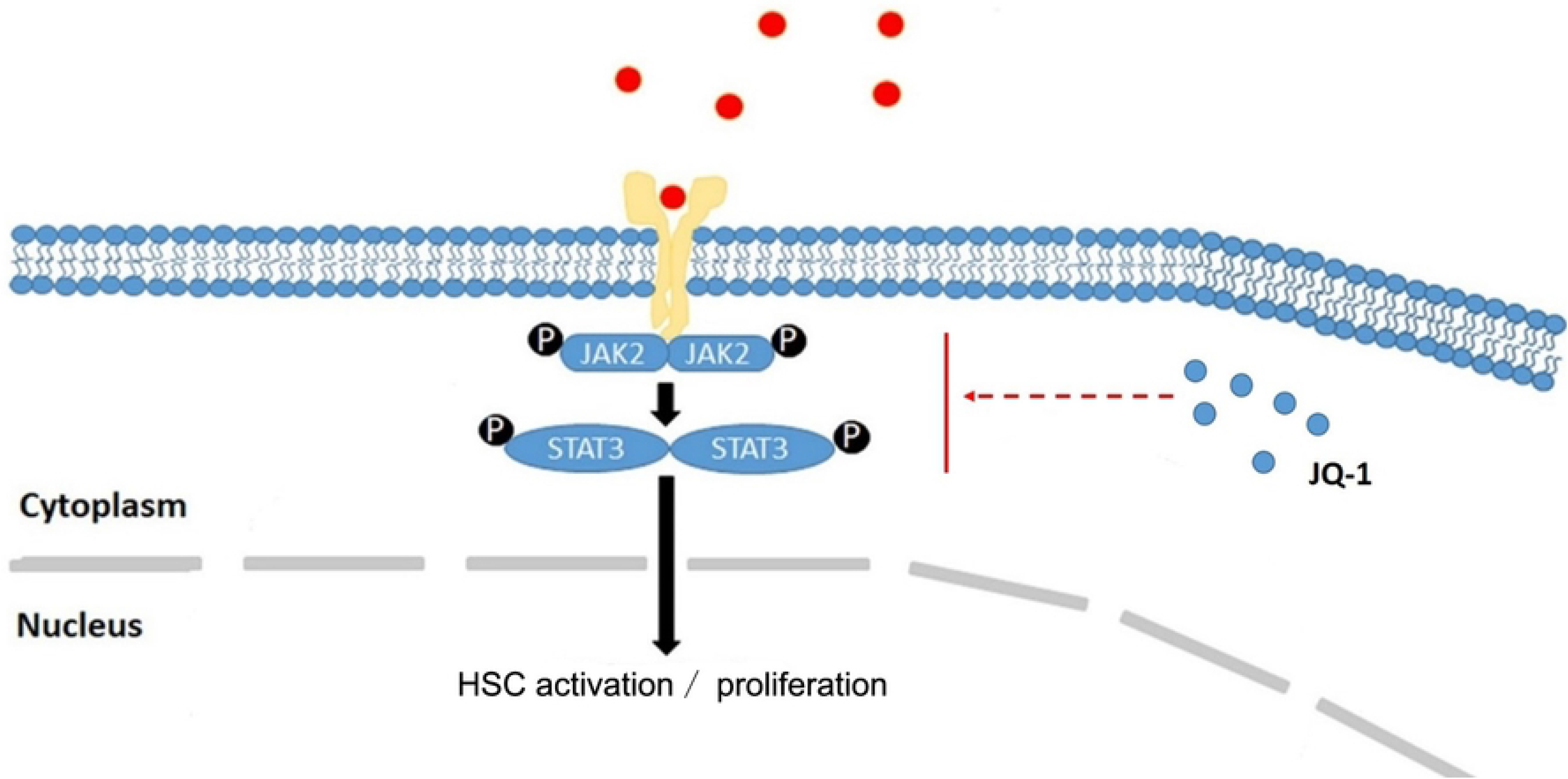
Schematic diagram of the inhibition of hepatic stellate cell (HSC) activation and proliferation by JQ-1. JQ-1 blocks downstream transcriptional regulation by binding to BRD4, thereby blocking phosphorylation of JAK2 and STAT3 in the JAK2/STAT3 signaling pathway and inhibiting HSC activation and proliferation.

In summary, the administration of JQ-1 decreased the degree of liver fibrosis caused by *S. japonicum*. JQ-1 inhibited the activation and proliferation of HSCs by blocking the phosphorylation of JAK2 and STAT3 in the JAK2/STAT3 signaling pathway to achieve this therapeutic effect. Our study provides new insight into the potential development of new therapeutic strategies for *Schistosoma*-induced liver fibrosis.

## Acknowledgments

We thank a native English speaker (Science/Medical English Editing for International Researchers) for modifying the manuscript.

## Data Availability

All relevant data are within the manuscript and its Supporting Information files.

